# Several FDA-approved drugs effectively inhibit SARS-CoV-2 infection *in vitro*

**DOI:** 10.1101/2020.06.05.135996

**Authors:** Hua-Long Xiong, Jia-Li Cao, Chen-Guang Shen, Jian Ma, Xiao-Yang Qiao, Tian-Shu Shi, Yang Yang, Sheng-Xiang Ge, Jun Zhang, Tian-Ying Zhang, Quan Yuan, Ning-Shao Xia

## Abstract

To identify drugs that are potentially used for the treatment of COVID-19, the potency of 1403 FDA-approved drugs were evaluated using a robust pseudovirus assay and the candidates were further confirmed by authentic SARS-CoV-2 assay. Four compounds, Clomiphene (citrate), Vortioxetine, Vortioxetine (hydrobromide) and Asenapine (hydrochloride), showed potent inhibitory effects in both pseudovirus and authentic virus assay. The combination of Clomiphene (citrate), Vortioxetine and Asenapine (hydrochloride) is much more potent than used alone, with IC50 of 0.34 μM.

## Main

As of 26 May 2020, the COVID-19 pandemic has claimed more than 342,000 lives, but yet effective drug is not available. It is time-consuming to develop vaccines or specific drugs for a disease caused by a novel defined virus like SARS-CoV-2. Re-purposing of approved drugs may be a faster way to find treatment for COVID-19. Verification of drugs that might suppress SARS-CoV-2 by prediction, including drugs against similar virus and broad-spectrum antiviral agents (BSAAs), is time-saving for drug re-purposing at the expense of missing some potential candidates. Integrative, antiviral drug repurposing methods based on big data analysis or molecular docking and molecular dynamics are time-saving and high throughput. However, drugs identified by virtual screening still need to be verified *in vitro* and *in vivo*.

In our previous research, a robust neutralization assay was established based on SARS-CoV-2 S-bearing vesicular stomatitis virus (VSV) pseudovirus and human ACE2-expressing BHK21 cells (BHK21-hACE2)^1^. Single-cycle infectious of recombinant VSV-SARS-CoV-2-Sdel18 mimics the entry of SARS-CoV-2. The infection of pseudovirus can be detected by fluorescence 12 hours after infection, enabling the assay time-saving for high-throughput screening^1^. This pseudovirus based assay is suitable for screening drugs that can block the infection of SARS-CoV-2.

In this study, the anti-SARS-Cov-2 potentiality of 1403 FDA approved drugs were quantitatively evaluated by the pseudovirus-based assay. The screening procedure was illustrated in Figure 1A and described in methods. The numbers of GFP-positive cells from drug treated wells were counted and divided by the number of infected cells from the well without treatment of drugs to calculate the relative value of infection rate. The results of two repetitions showed that most of drugs did not inhibit viral infection (Figure 1B). Forty-four drugs with relatively better inhibitory effect were selected for further validation. In the second round of screening, inhibition of VSV-SARS-CoV-2-Sdel18 virus infection and cell cytotoxicity were both detected (Supplementary Figure 1). Among them, 31 drugs were excluded due to cytotoxicity. 13 drugs were selected for analysis of specificity to VSV-SARS-CoV-2-Sdel18 and verification by authentic SARS-CoV-2 assay.

**Figure 1:**
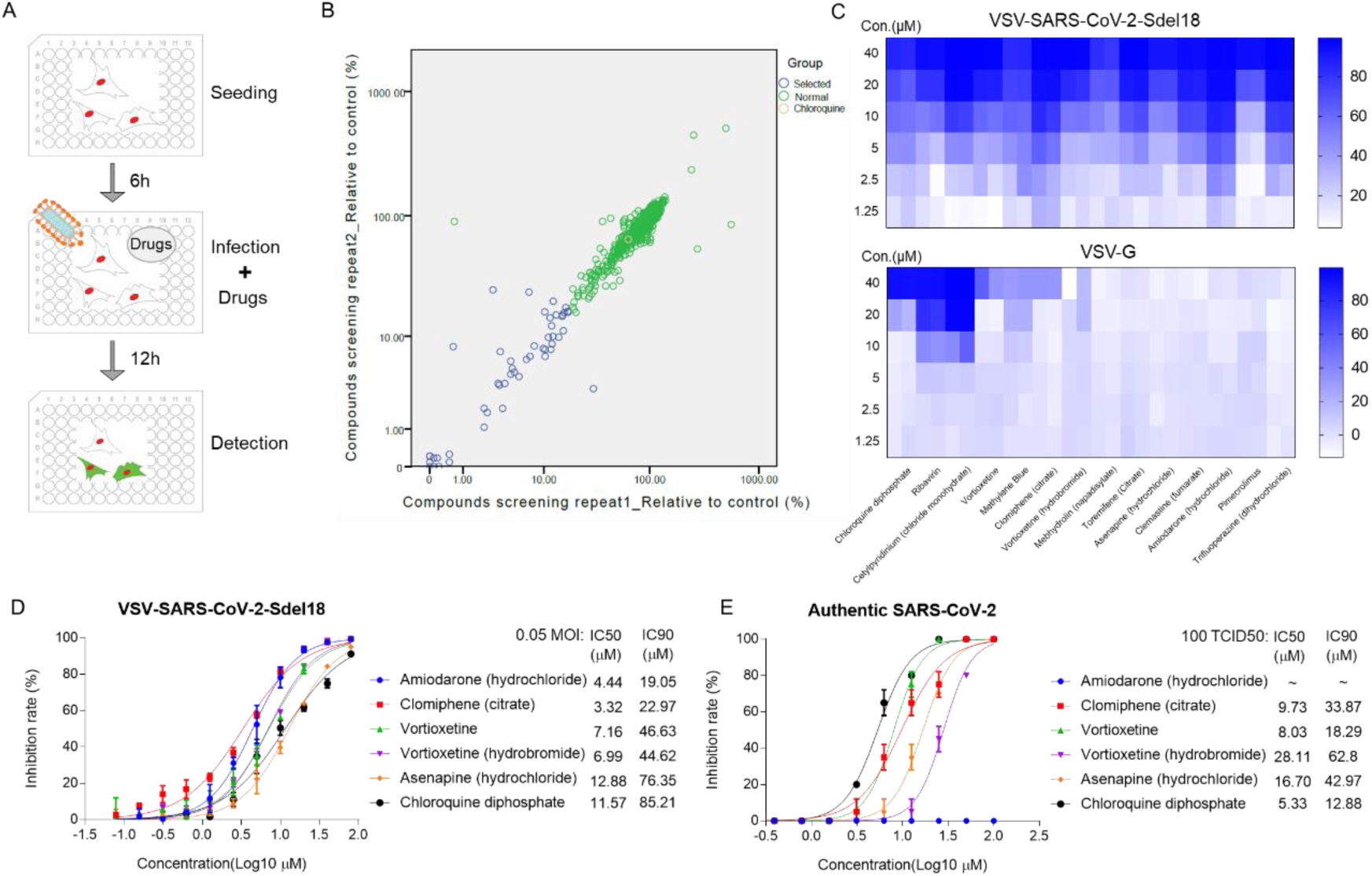
Re-purposing of FDA-approved drugs for inhibiting SARS-CoV-2 infection. A Schematic diagram of the screening process of the pseudovirus model. B The first round of screening for compounds could inhibit the infection of VSV-SARS-CoV-2-Sdel18. The abscissa and ordinate respectively indicate two repetitions of screening. Yellow circle: chloroquine. Blue circles: 44 drugs selected for the second round of screening, Green circles: remaing FDA-approved drugs. C Screening for the compounds specially block the spike protein of SARS-CoV-2 mediated viral entry. The inhibitory effect to VSV-SARS-CoV-2-Sdel18 and VSV-G of 13 seleted compounds was evaulated to exclude drugs inhibit infection or expression of VSV-G. Colorbar indicates inhibiton rate. Analyze the inhibitory effect of five selected compounds in VSV-SARS-CoV-2-Sdel18 pseudovirus assay (D) and authentic SARS-CoV-2 assay (E).

To verify whether these selected drugs act on spike protein of SARS-CoV-2 on the pseudovirus or the VSV backbone, we evaluated the inhibitory effect of these compounds on VSV-G (The sequence of GFP was inserted into the genome of VSV, so that the infection of VSV could be indicated by green fluorescence.). There were three compounds, including chloroquine diphosphate, ribavirin, and cetylpyridinium (chloride monohydrate), exhibited significant inhibitor effects on VSV-G, whereas no effect was noted for other compounds (Figure 1C).

Considering the inhibitory effect and cytotoxicity (Supplementary Figure 2), five compounds inhibited the infection of VSV-SARS-CoV-2-Sdel18 pseudovirus specifically, including Clomiphene (citrate), Amiodarone (hydrochloride), Vortioxetine, Vortioxetine (hydrobromide) and Asenapine (hydrochloride), were selected and the function of these compounds was confirmed using authentic SARS-CoV-2 assay (Figure 1D and E). Among them, the inhibitory effects of Clomiphene (citrate) and Vortioxetine were comparable to Chloroquine diphosphate, while Vortioxetine (hydrobromide) and Asenapine (hydrochloride) were slightly less effective. Whereas Amiodarone (hydrochloride) inhibited the infection of pseudovirus efficiently with IC50 around 4.44 μM, but it showed no effect on authentic SARS-CoV-2 virus infection even used at a concentration of 100 μM.

To further evaluate the potential of applying these drugs in prophylaxis and combination therapy, we treated the cell with pseudovirus and different drug combinations. The drug combinations were added either at the same time of pseudovirus infection or 6 hours pre-infection (Table 1 and Supplementary Figure 3). The combination of Clomifene (citrate), Vortioxetine and Asenapine (hydrochloride) showed best effect when used both at the time of infection and pre-infection, with IC50 about 1.93 μM and 0.34 μM respectively. The combination of Clomifene (citrate) and Vortioxetine had a comparable effect, with IC50 about 2.36 μM and 0.69 μM respectively. The combination of drugs decreases the concentration of each drug required to block virus infection, which may reduce the side effects of drugs. However, it still remains to be evaluated whether these drugs can be used together *in vivo*.

**Table 1.** The inhibitory potency of combination of drugs

| Drug | Pre-treatment |  |  | Co-treatment |  |  |
| --- | --- | --- | --- | --- | --- | --- |
|  | IC50(μM) | IC90(μM) | CC50(μM) | IC50(μM) | IC90(μM) | CC50(μM) |
| Clo+Vor | 0.69 | 4.81 | 14.47 | 2.36 | 11.80 | 23.55 |
| Clo+Ase | 1.60 | 10.39 | 21.39 | 3.71 | 20.06 | 36.20 |
| Vor+Ase | 2.08 | 11.06 | 14.12 | 3.52 | 17.47 | 28.00 |
| Clo+Vor+Ase | 0.34 | 5.01 | 14.67 | 1.93 | 9.42 | 16.83 |
| Clo+CQ | 2.57 | 12.81 | 28.01 | 7.32 | 17.99 | 37.98 |
| Vor+CQ | 3.03 | 13.00 | ~ 18.93 | 5.39 | 17.87 | 25.69 |
| Ase+CQ | 5.73 | 55.30 | 128.00 | 12.70 | 58.87 | na |
| Clo | 2.94 | 10.16 | 24.94 | 9.53 | 19.57 | 54.35 |
| Vor | 3.00 | 13.83 | 28.53 | 6.77 | 22.28 | 27.05 |
| Ase | 17.69 | 127.10 | na | 28.13 | 117.60 | na |
| CQ | 9.27 | 35.22 | 349.10 | 27.60 | 106.20 | na |

“Clo” means Clomiphene (citrate), “Vor” means Vortioxetine, “Ase” means Asenapine
(hydrochloride) and “CQ” means Chloroquine diphosphate.
“Pre-treatment” means cell was treated with drugs 6 hours before infection, while “Co-treatment” means cells were treated with drugs at the time of infection.
IC50, IC90 and CC50 were calculated using prism software (GraphPad). “na” means the value can’t be calculated. MOI=0.1

There are currently no approved drugs or vaccines against SARS-CoV-2. Hundreds of clinical trials have been initiated to establish evidence around investigational drugs and vaccine candidates. Several approved or on-trial drugs are under investigation for treating COVID-19 patients, including hydroxychloroquine, arbidol, remdesivir, favipiravir, lopinavir/ritonavir, and others. However, it seems that the therapeutic effects of these drugs are not ideal according to the published clinical data. Only a multicenter randomized open-label phase 2 trial showed that a triple combination of interferon beta-1b (inject), lopinavir/ritonavir and ribavirin (oral take) is demonstrated to be effective in suppressing the shedding of SARS-CoV-2, when given within 7 days of symptom onset^2^. Re-purposing of other approved drugs may bring more candidates for the treatment of COVID-19.

Drugs that block the infection of SARS-CoV-2 may alleviate disease progression, protect health care workers and other populations at high risk of infection. In this research, an efficient VSV-SARS-CoV-2-Sdel18 pseudovirus model was applied to identify candidates that can inhibit infection of SARS-CoV-2 from 1403 approved drugs. Five drugs, which haven’t been identified before, showed comparable or superior inhibitory effect to chloroquine in this model. The effect was also confirmed using authentic SARS-CoV-2 assay and four of them can also inhibit the infection of authentic SARS-CoV-2 virus.

Clomifene Citrate is a selective estrogen receptor modulator and a non-steroidal fertility medicine. It has a long history of use since 1967 and has the advantages of oral availability, good safety, and tolerability profiles. Johansen et al. identified Clomiphene as potent inhibitors of Ebola virus infection^3^. It may inhibit Ebola virus through inducing accumulation of cholesterol in endosomal compartments and blocking the release of viral genome to cytoplasm^3-5^. The viral entry of SARS-CoV-2 includes the endocytosis of enveloped viral particle, priming of spike protein by protease, fusion between viral and cellular membranes and release of viral genome, which is similar to Ebola virus ^6-8^. Therefore, the Clomiphene may impair SARS-CoV-2 infection via the same pathway as Ebola virus.

Vortioxetine is an antidepressant drug that is used to treat major depressive disorder in adults. Vortioxetine was safe and well tolerated, it was approved in 2013 ^9^. So far, no previous study described its antiviral roles. It is reported that sever COVID-19 patients have a high probability of suffering from mental illness. Recently, another antidepressant drug fluvoxamine is evaluated for the potential to treat COVID-19 by researchers from the Washington University School of Medicine, because the drug may prevent an overreaction of the immune system called cytokine storms, which could result in life-threatening organ failure. The antiviral mechanism of Vortioxetine remains unknow. However, it may bring physical and psychological benefits for COVID-19 patients.

Asenapine is an antipsychotic medicine that is used to treat schizophrenia and bipolar I disorder and has been approved since 2009. Notably, it showed less cytotoxicity in this study comparing to other drugs that could inhibit the infection of SARS-CoV-2.

In summary, our study identified four FDA-approved drugs that have the potential to suppress SARS-CoV-2 infection. The robust assay based on VSV-SARS-CoV-2-Sdel18 pseudovirus screened out the potential drugs with high efficiency, then the inhibitory effect was confirmed by authentic SARS-CoV-2 assay. The inhibitory effect of Vortioxetine and Clomifene is superior and the mechanism of these drugs seems different from Chloroquine. The combination of Clomifene (citrate), Vortioxetine and Asenapine (hydrochloride) greatly decreases the IC50/IC90 of blocking virus infection. The clinical safety of these compounds has been evaluated and the availability of pharmacological data are expected to enable rapid preclinical and clinical evaluation for treatment of COVID-19. Based on the existing clinical results, it seems that it is difficult for one particular drug alone to significantly benefit COVID-19 patients, and combination therapy is more likely to make the patient recover faster. This work identified novel drugs that suppress the infection of virus and provided more candidates for post-exposure prophylaxis and combination therapies. Notice that no test *in vivo* has been conducted and the mechanism of these compounds also remains unknown. More researches are required to support the clinical application of these drugs for treatment of COVID-19.

## Materials and Methods

### Cells and samples

Vero-E6 (American Type Culture Collection [ATCC], CRL-1586), Vero (ATCC, CCL-81), BHK21-hACE2^1^ cells were maintained in high glucose DMEM (SIGMA-ALDRICH) supplemented with 10% FBS (GIBCO), penicillin (100 IU/mL), streptomycin (100 μg/mL) in a 5% CO_2_ environment at 37°C and passaged every 2 days. In addition, the culture medium of BHK21-hACE2 contains puromycin (2 μg/mL). The FDA-approved drug library, including 1403 compounds (10 mM DMSO solutions), and Chloroquine diphosphate were bought from MedChemExpress (MCE).

### Pseudovirus-based assay

VSV pseudovirus carrying truncated spike protein of SARS-CoV-2, named VSV-SARS-CoV-2-Sdel18 virus, was packaged as previously described^1^. VSV-G was prepared in similar way ^10^. In the first round of screening, all compounds were diluted to 20 μM and mixed with VSV-SARS-CoV-2-Sdel18 virus, the volume of diluted compounds and virus are 80 μL and 20 μL respectively. Added 80 μL final mixture, which containing compounds (16 μM) and pseudovirus (MOI=0.05), to pre-seeded BHK21-hACE2. After 12h incubation, fluorescence images were obtained by ImmunSpot@S5 UV Analyzer (Cellular Technology Limited) or Operetta CLS (PerkinElmer). For quantitative determination, the numbers of GFP-positive cell for each well were counted to represent infection performance. The reduction (%) in GFP-positive cell numbers was calculated to show the inhibitory effect of compounds. In the second round of screening, selected compounds were diluted to 50 μM, then serial two-fold dilutions are used to prepare diluted analytes. 80 μL diluted compounds were mixed with 20 μL VSV-SARS-CoV-2-Sdel18 or VSV-G and the mixture were added to pre-seeded BHK21-hACE2. The results were obtained as described previously. To analysis the IC50 of selected compounds, the compounds were diluted to 100 μM as the first work concentration and 0.098 μM as the smallest concentration. Still mixed 80 μL diluted compounds with 20 μL VSV-SARS-CoV-2-Sdel18 virus. The remaining procedures were same as previous assay. The cytotoxicity of compounds was analyzed by Cell Counting Kit-8 (CCK-8, MCE). To evaluate the effect of drug combinations, the drugs were also diluted to 100 μM (the concentration of each drugs is 100 μM in mixture) and prepared serious dilutions. To evaluate the potential of applying these drugs in prophylaxis, the cell was pre-treated with 80 μL diluted drugs, 6 hours later, add 20 μL virus to the culture medium (MOI=0.1).

### Authentic SARS-CoV-2-based assay

Vero cells were seeded 24 hours before the infection in a 96-well plate (Costar). On the day of infection, the cells were washed twice with PBS. Candidate drugs were diluted 2-fold seriously by medium supplemented with 2% FBS (GIBCO), penicillin (100 IU/mL), streptomycin (100 μg/mL). Aliquots (40 μL) of diluted drugs (200 μM as initial concentration) was added to 40 μL of cell culture medium containing 100 times the tissue culture infective dose (TCID50) of the BetaCoV/Shenzhen/SZTH-003/2020 strain virus (GISAID access number: EPI_ISL_406594) on a 96-well plate in decuplicate and incubated at 37 °C for 2 hours in CO_2_ 5% vol/vol. After incubation, virus drugs mix was then added to cells in 96-well plates and plates were incubated at 37 °C with microscopic examination for cytopathic effect after a 5-day incubation. The complete absence of cytopathic effect in an individual culture well was defined as protection. The values of IC50 were calculated using prism software (GraphPad).

### Statistic

The relative value or inhibition rate of candidate drugs were calculated according to the decrease of GFP positive cell number (for pseudovirus-based assay) or cytopathic effect (for authentic SARS-CoV-2-based assay). The IC50 (the half maximal inhibitory concentration), IC90 (the concentration for the 90% of the maximum inhibition) and CC50 (the 50% cytotoxic concentration) values were calculated with non-linear regression, i.e. log (inhibitor) vs. normalized response – Variable slope or log(agonist) vs. response—Find ECanything using GraphPad Prism 7.00 (GraphPad Software, Inc., San Diego, CA, USA).

## Author Contribution

Hua-Long Xiong, Tian-Ying Zhang, Quan Yuan and Ning-Shao Xia had full access to all of the data in the study and take responsibility for the integrity of the data and the accuracy of the data analysis. Study concept and design: Tian-Ying Zhang, Quan Yuan and Ning-Shao Xia. Acquisition of data: Hua-Long Xiong, Jia-Li Cao, Jian Ma, Xiao-Yang Qiao, Tian-Shu Shi. Chen-Guang Shen and Yang Yang performed the authentic virus assay. Analysis and interpretation of data: Hua-Long Xiong, Jia-Li Cao, Tian-Ying Zhang. Drafting of the manuscript: Jia-li Cao, Tian-Ying Zhang, Hua-Long Xiong. Critical revision of the manuscript for important intellectual content: Sheng-Xiang Ge, Jun Zhang. Study supervision: Tian-Ying Zhang, Quan Yuan and Ning-Shao Xia.

## Acknowledgements

This work was supported by the National Natural Science Foundation of China (Major Program: 81993149041, 81702006), Science and Technology Major Project of the Fujian Province (2020YZ014001), Xiamen Science and Technology Major Project (3502Z2020YJ02), Shenzhen Science and Technology Innovation Commission for Research and Development Project (Grants JCYJ20190809183205622).

