## Supplementary file for "Several FDA-approved drugs effectively inhibit SARS-CoV-2 infection *in vitro*"

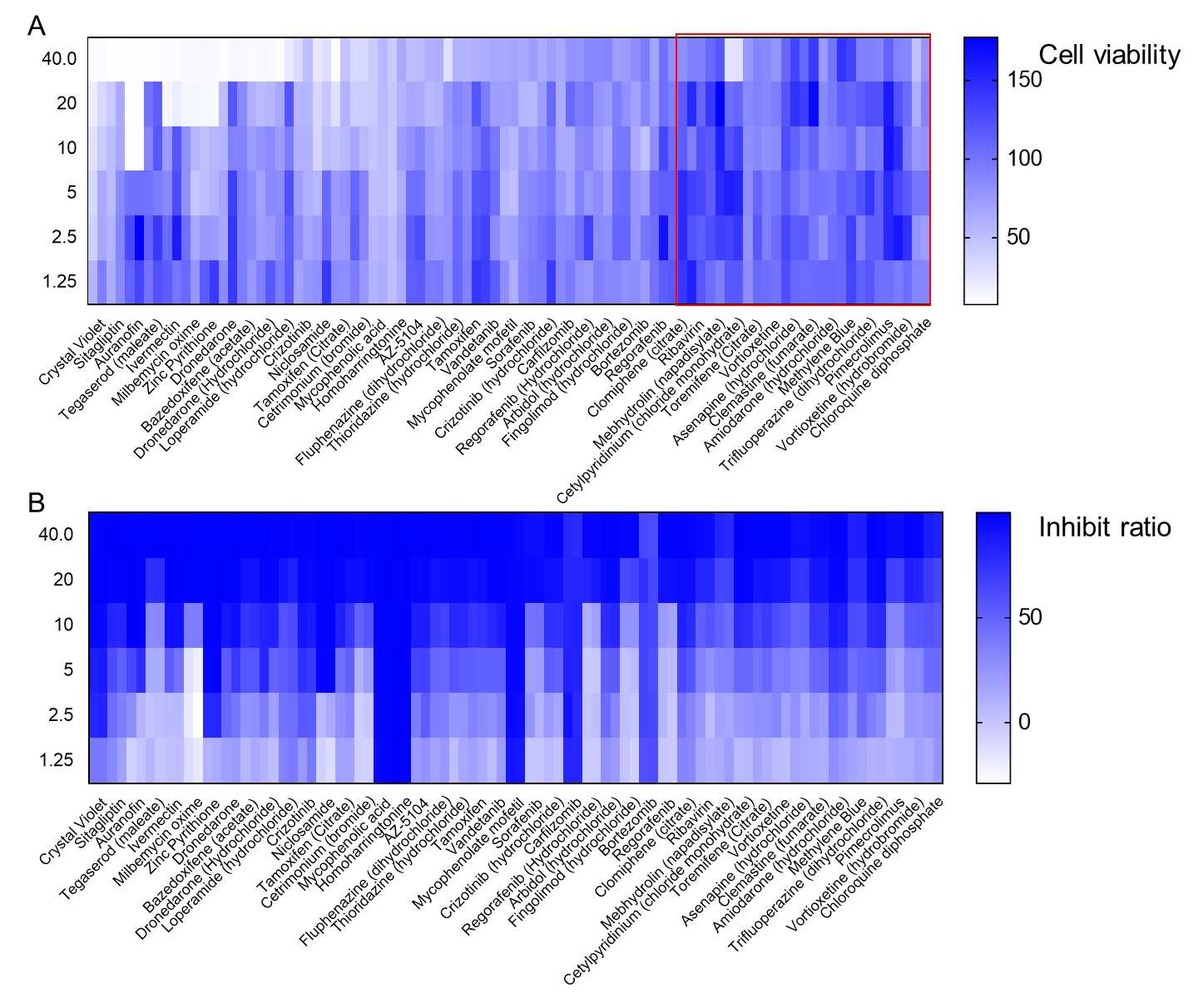

Supplementary Figure 1: The second round of screening for compounds could inhibit the infection of VSV-SARS-CoV-2-Sdel18. 44 drugs were selected according to the result of first rounds of screening. The inhibitory effect and cytotoxicity of these drugs were further evaluated by serious dilution. A The cell viability analyzed by CCK-8. B The inhibitory effect to the infection of VSV-SARS-CoV-2-Sdel18. Ordinate stands for the concentration of drugs (μM).

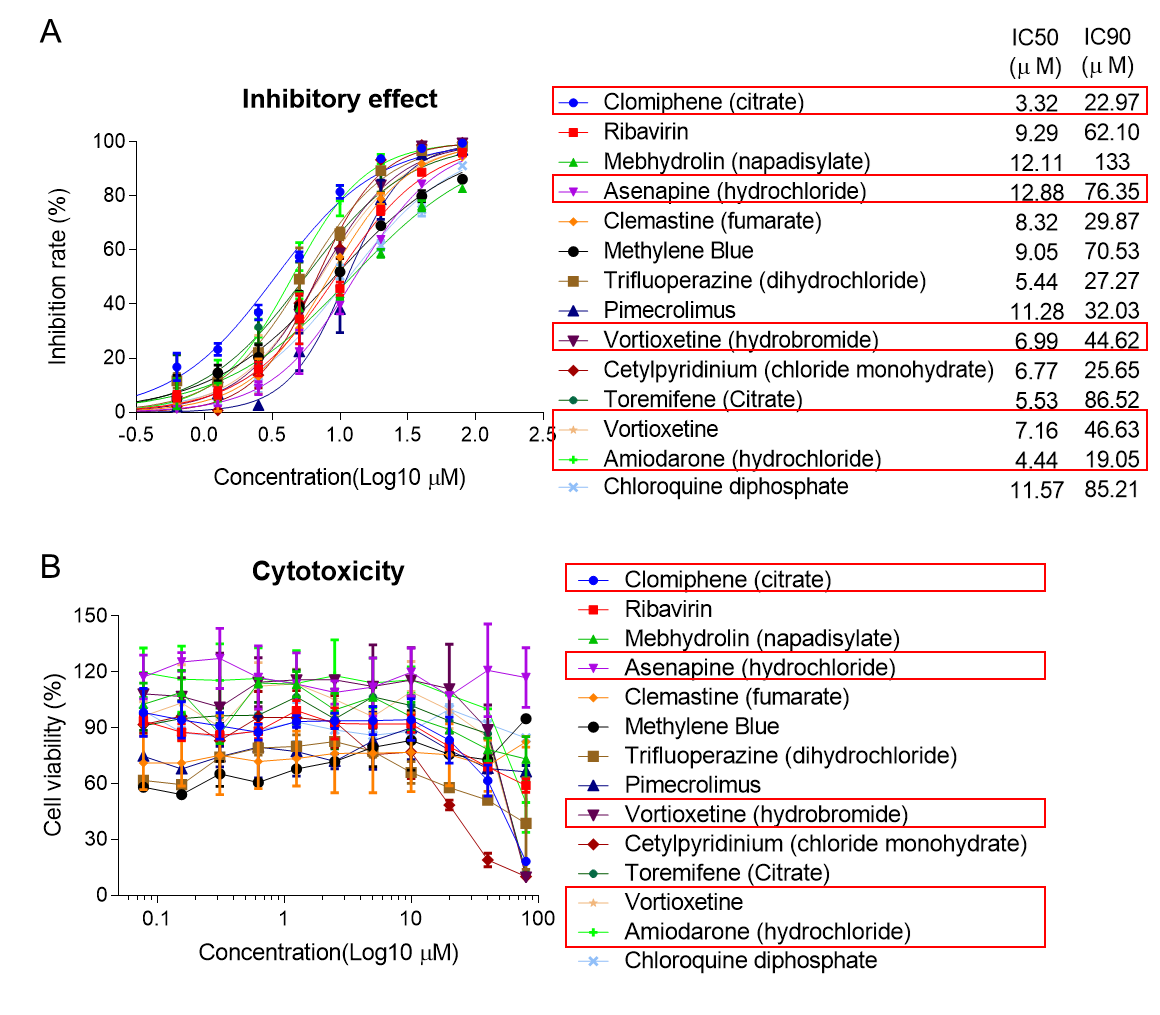

Supplementary Figure 2: Further analysis of the inhibitory and safety of 13 selected drugs. A The 13 selected drugs were seriously diluted and analyzed the IC50 and IC90 to VSV-SARS-CoV-2-Sdel18 (MOI=0.05). B Cytotoxicity analysis results of the corresponding concentration of drugs.

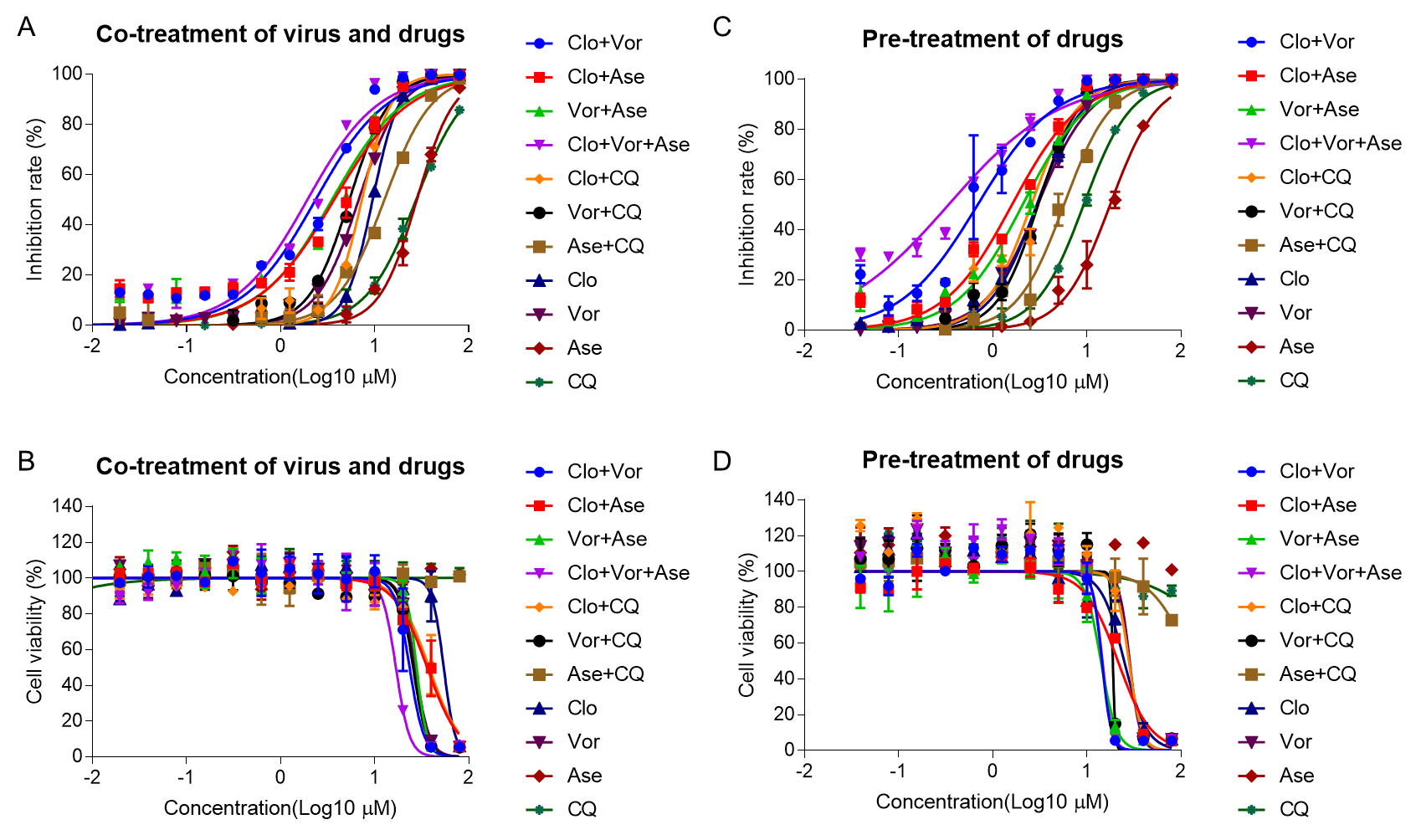

Supplementary Figure 3: Evaluate the inhibitory effect and cytotoxicity of drug combinations. A The serious dilutions of different drug candidates and drug combinations were prepared and mixed with VSV-SARS-CoV-2-Sdel18 virus (MOI=0.1), the mixture was added to pre-seeded BHK21-hACE2 cells. The fluorescence of cell was detected to analyze the inhibitory effect and the cytotoxicity was analyzed by CCK-8 (B). C To evaluate the function of these drugs in prophylaxis, BHK21-hACE2 cells were pre-treated with seriously diluted drug candidates or drug combinations, 6 hours later, added VSV-SARS-CoV-2-Sdel18 virus to infect cells (MOI=0.1). The fluorescence of cell was detected to analyze the inhibitory effect and the cytotoxicity was analyzed by CCK-8 (D).
